# A DNA foundation model predicts osteoporosis risk genes without proximity bias

**DOI:** 10.64898/2026.03.09.707383

**Authors:** Cristian Regep, Chantriolnt-Andreas Kapourani, Emre Sofyali, Agnieszka Dobrowolska, Giannis Loukas, Andrew Anighoro, Eleonora Canale, Torsten Gross, Marco P. Licciardello, Revant Gupta, Sorina Maciuca, Tariq Desai, Alice Del Vecchio, Cameron Field, Karl Gemayel, Avelino Javer, Zicong Zhang, Rika Tsujikawa, Fumitaka Inoue, Edith. M. Hessel, Jake P. Taylor-King, John Whittaker, David Roblin, Rebecca E. McIntyre, Lindsay M. Edwards

**Affiliations:** Relation Therapeutics, London, UK; MRC Biostatistics Unit, University of Cambridge; Institute for the Advanced Study of Human Biology (WPI-ASHBi), Kyoto University, Kyoto, Japan; The Francis Crick Institute, London, UK

**Author notes:** joint senior authors (Rebecca E. McIntyre, Lindsay M. Edwards).

## Abstract

Targets supported by human genetic associations are more than twice as likely to progress from clinical development to approval. Genome-wide association studies are the largest source of genetic evidence for disease risk but linking non-coding variants to effector genes remains a significant barrier to identifying causal targets. Current gene-mapping approaches suffer from proximity bias, largely ignoring distal genes. Here we introduce Rosalind, a DNA foundation model fine-tuned on human genetic variation from GTEx, that directly predicts variant-gene regulatory relationships from sequence without relying on nearest-gene heuristics. We demonstrate Rosalind’s accuracy through extensive benchmarking, apply it to multiple complex traits to establish broad utility, and provide experimental validation in osteoporosis using a translational osteoblast assay. We demonstrate that genes distal to osteoporosis risk variants were significantly more likely to alter a bone formation phenotype than nearest genes. Together, these results highlight deep learning-based regulatory models as a general and scalable framework for translating novel genetic insights to drug discovery.

## Introduction

Clinical drug development continues to face high attrition rates, with approximately 90% of candidates failing to reach the market, making evidence-based strategies to improve success rates extremely valuable. One of the most robust predictors of success is human genetic evidence [1, 2]. Specifically, if a gene linked to a candidate drug target is statistically associated with a phenotype relevant to the therapeutic indication, the probability of success more than doubles. This effect is primarily thought to arise because genetic linkage infers disease causality not correlation, and genetics is not confounded by the limits of traditional epidemiological approaches [4], or reverse causation seen in differential expression studies [3]. As a result, integrating human genetics into target identification pipelines increases the likelihood of nominating biologically grounded and therapeutically relevant drug targets.

Early efforts to integrate human genetic evidence into the drug development process overwhelmingly prioritized genes implicated in monogenic disorders, or variants with unambiguous gene-phenotype relationships and high penetrance [5]. However, the most abundant source of genetic variation associated with disease, or traits, comes from genome-wide association studies (GWAS). Approximately 90% of all the GWAS signals fall in non-coding regions, implicating regulatory mechanisms rather than coding variation [6]. Establishing the regulatory mechanism whereby a variant (or variants) influence physiology, with concomitant increases in an individual’s risk of disease, remains challenging. While variant-to-gene (V2G) methods exist to map non-coding variants to their effector gene (eGenes) [7, 8], the majority rely either implicitly or explicitly on the assumption that the nearest gene (in terms of base pair distance) is most likely to be causal [9] This is overly simplistic as linear base pair distance does not necessarily reflect physical closeness in the nucleus, where DNA folds and packs tightly in three dimensional space [10]. There is significant variation across loci, for example in a study of variants associated with bone mineral density, we found that the number of genes varied from 0 to 30 (Fig. 3b). At present, prevailing approaches either assign the nearest gene as the putative effector or rely on integrative approaches like locus to gene (L2G, Open Targets;11]) that incorporate orthogonal ‘omic datasets. However, these datasets often provide little additional information on top of distance, and predictions remain strongly driven by gene proximity [12] (Fig. 2a). Intuitively, this will work best in relatively simple situations (for example where only one gene is within some distance of a variant), but will be unreliable in complex regions with many plausible genes in close proximity. Recently, colocalisation of single cell expression quantitative trait loci (sc-eQTL) maps with GWAS variants has demonstrated enrichment for disease-related eGenes at distal enhancers [23]. While this approach is powerful, widespread adoption is hampered by the requirement to process hundreds of clinically accessible biopsies, which is operationally challenging and resource intensive.

Recent advances in sequence-based deep learning models provide an opportunity to address the distal V2G challenge in any tissue and using existing datasets. Transformer-based architectures, such as Enformer [15], Borzoi[29] and AlphaGenome[30] have demonstrated the ability to capture both regulatory syntax and long-range enhancer-promoter interactions from DNA sequence alone, without explicit supervision. However, these foundation models are typically trained on the reference genome and face challenges in generalising to individual-level genetic variation or predicting genotype-specific expression [14]. Moreover, their utility to improve causal gene discovery across diverse disease contexts hasn’t been fully explored. Here we introduce Rosalind, a DNA foundation model fine-tuned on GTEx expression eQTL data to learn how genetic variation alters gene regulation. We benchmark its performance against comparable approaches, demonstrating consistently improved accuracy in identifying distal causal variant-gene pairs and enrichment for biologically plausible distal eGenes that overlap with known drug targets.

Using osteoporosis as a translational case study, we used CRISPR/Cas9 to knock-out Rosalind predicted disease risk genes in a human osteoporosis-relevant assay of osteoblast activity. Osteoblasts are one of the key effector cell types of osteoporosis and in vitro, they deposit quantifiable levels of hydroxyapatite, a major component of bone mineral. We demonstrate that Rosalind’s distal predictions are significantly enriched as hits in the assay versus nearest controls, identifying functional links overlooked by nearest-gene heuristics and highlighting a novel role for genes involved in maintenance of primary cilia structure in genetic predisposition to low bone density and risk of osteoporosis. Collectively, these findings highlight the potential of sequence-based foundation models to overcome proximity bias and provide a scalable framework for causal gene discovery.

## Results

### Capturing regulatory logic across diverse sequence contexts and scales

A central challenge in regulatory genomics is that biological signals operate across multiple spatial scales, from local promoter grammar to long-range enhancer-promoter interactions. Transformer-based architectures such as Enformer [15], Borzoi[29] and AlphaGenome[30] provide strong performance on large genomic windows by integrating distal regulatory information. Conversely, architectures like GeneGenie [17] are designed for high-resolution modelling of promoter-proximal logic, explicitly encoding structural constraints and strand interactions to capture local regulatory syntax.

While transformers offer expansive receptive fields and flexible attention-based representations, it is not yet clear whether mechanisms engineered for long-range integration inherently encode the inductive biases needed for fine-scale promoter grammar in order to capture the effects of individual variants. Even when normalized to a comparable receptive field, the Enformer [15] architecture diverges fundamentally in its structural logic by employing relative positional basis functions to model interactions as distance-dependent decay and utilizing attention pooling to selectively weight specific sequence features. At the other extreme, the local architecture proposed by GeneGenie [17] enforces sequential and structural constraints, utilizing a bidirectional LSTM downstream to capture grammatical dependencies, and specialized 2D convolutions to explicitly model forward-reverse strand interactions, rather than learning them implicitly through data augmentation as Enformer [16].

To empirically resolve these divergent architectural philosophies, we benchmarked both approaches on two diverse MPRA datasets: a high-throughput yeast promoter screen (Fig. 1a) and an independent osteoblast enhancer assay (Fig.1 b.). Across both contexts, we observe that the Enformer-style architecture achieves strong predictive performance, suggesting that long-range transformer architectures can in practice scale down to capture fine-grained regulatory syntax, despite their original design goals.

**Fig. 1.**
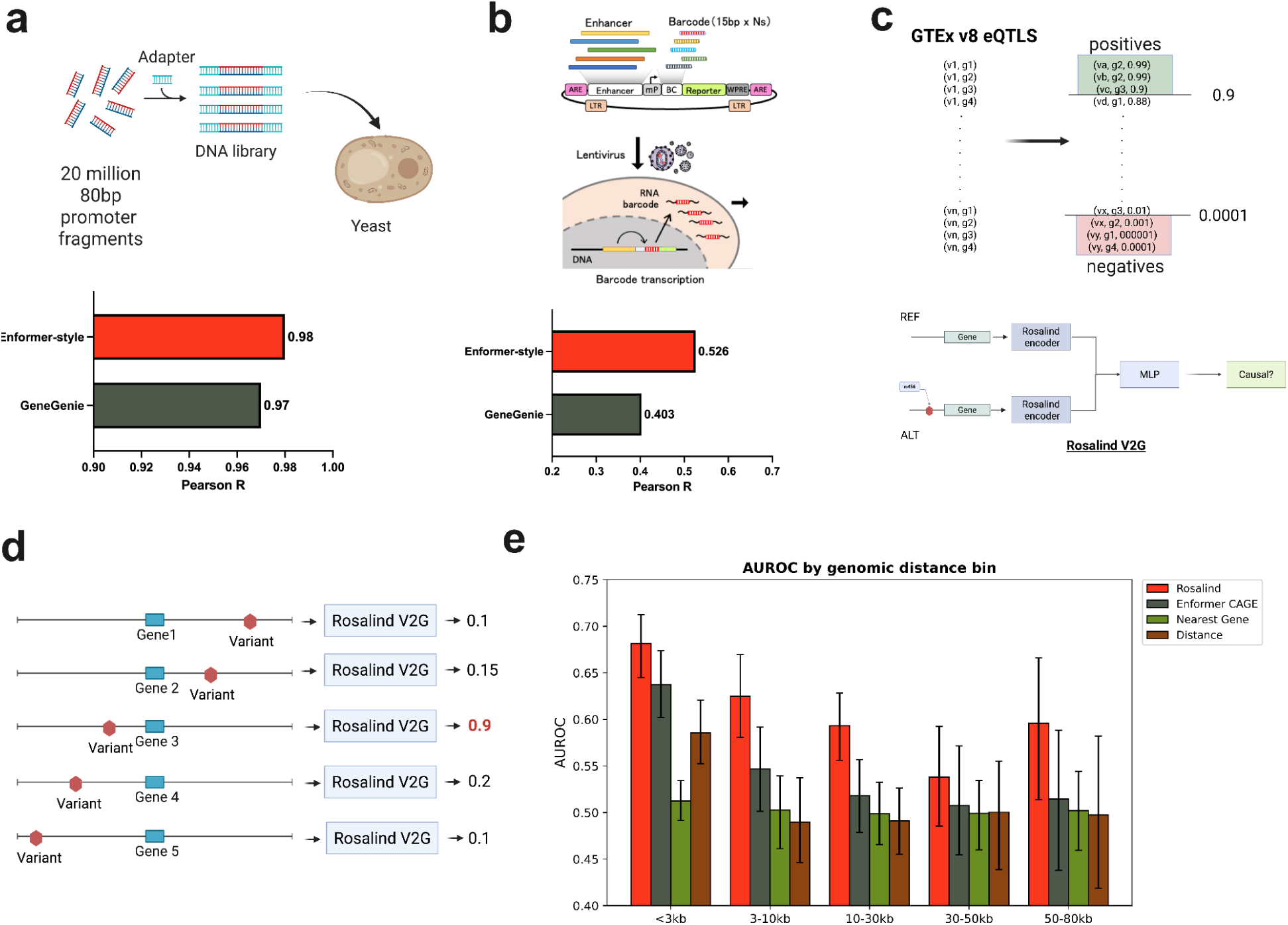
Rosalind architecture. a. Enformer-style architecture benchmarked for local-scale regulatory prediction on random yeast promoters with in vivo fitness data and the evaluation set up from GeneGenie [17] b. Same set-up from a. applied to enhancer saturation mutagenesis in primary human osteoblasts c. Fine-tuning set-up for causal variant-gene pairs using fine-mapped GTEx eQTLs. Variants were labeled positive/negative using PIP thresholds [19] d. Rosalind can be used to score all genes within a set receptive field, estimating the likelihood that an index variant alters their expression, enabling systematic causal-gene prioritization. e. AUROC for causal gene-prediction across different models, stratified by the distance to the causal gene

### Rosalind learns variant-to-gene links from fine-mapped cis-eQTLs

Building on these results we created Rosalind, based on an Enformer-style architecture where the relative positional encodings are replaced with Rotary Positional Embeddings (RoPE) to enhance long-range distance encoding and, crucially, we also fine-tune the foundation model as part of a separate causality binary classification task. We used a dataset of approximately 17,000 variant–gene pairs derived from fine-mapped GTEx cis eQTLs [18]. Fine-mapped signals were aggregated into a tissue-agnostic framework, and variant–gene pairs were labelled using established posterior inclusion probability (PIP) thresholds: variants with PIP > 0.9 were designated as high-confidence positives, whereas those with PIP < 0.01 served as negatives (Fig. 1c). For each variant-gene pair, Rosalind encodes both the reference and alternate alleles within a window and produces allele-specific embeddings for each. Then a lightweight multi-layer perceptron takes these embeddings and learns, via cross-entropy loss, to discriminate causal variants from non-causal ones relative to the reference sequence (Fig. 1c). During fine-tuning, the weights of the underlying DNA encoder remained unfrozen, allowing the Rosalind model to directly refine its representation of regulatory interactions in the presence of allelic perturbations. The resulting model enables assessing the impact of individual variants on its neighbouring genes (Fig. 1d).

### Benchmarking Rosalind against a foundation model in a tissue-agnostic setting

To evaluate whether finetuning Rosalind leads to improvement in identifying causal V2G pairs; we benchmarked it on a held-out set derived from GTEx fine-mapped cis-eQTLs. We compared Rosalind with Enformer (Fig. 1e), which was adapted to the same prediction task by aggregating the change in predicted CAGE signal between the reference and alternate alleles (Methods). We also included two baseline predictors based on genomic distance to TSS or whether the linked gene is the nearest gene to evaluate whether Rosalind can avoid proximity-based mis-assigments. While performance declines with increasing genomic distance, reflecting greater uncertainty in V2G links at longer distances, Rosalind generally outperforms Enformer-CAGE and baseline predictors.

### Rosalind generalises across diverse GWAS traits

We next asked whether Rosalind could be applied broadly to identify likely causal genes across complex traits and diseases, beyond the GTEx-derived eQTL setting used for model fine-tuning. To assess generalisability, we applied Rosalind to several large-scale GWAS datasets spanning diverse biological domains: type 2 diabetes, hypertension, asthma and psoriasis. These disease traits were selected due to having the highest number of genes with approved drugs in ChEMBL, a gold standard data source that is known to have less proximity bias than manually curated disease-gene pairs [11]. For each trait, we scored all genes within 131kbp of lead variants and identified high-confidence effector gene candidates.

Across all traits evaluated, Rosalind prioritised substantially more distal genes (i.e. not the nearest gene to a lead variant) compared to the L2G model in Open Targets. Specifically, Rosalind produced gene-sets with more than two-fold higher proportions of distal genes (Fig. 2b), whereas L2G predictions across therapeutic areas are dominated by nearest gene assignments and typically contain <20% distal genes (Fig. 2 a,b).

**Fig. 2.**
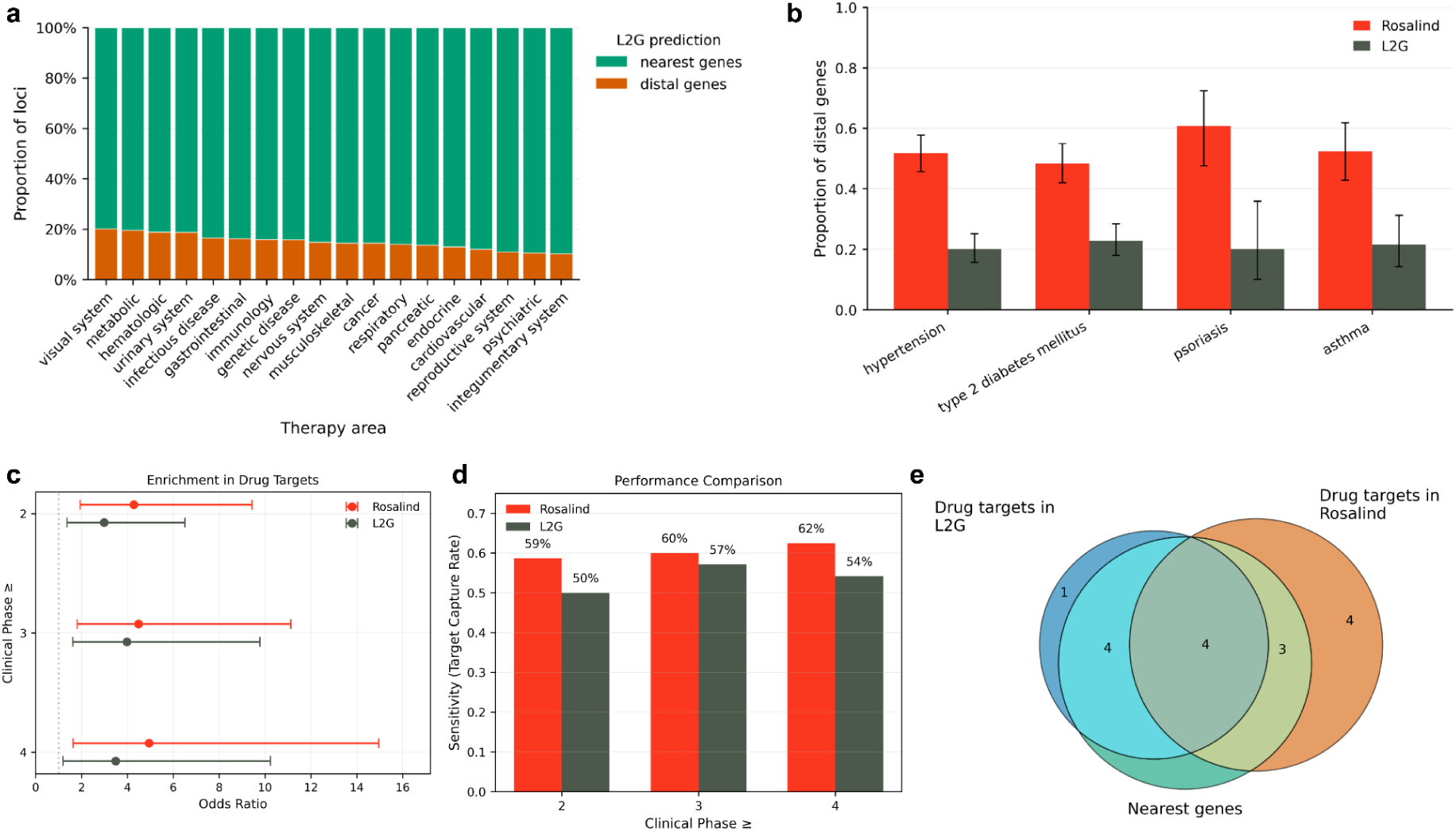
Benchmarking Rosalind on GWAS loci and clinically validated drug targets from OpenTargets+. a. Proportion of L2G predictions assigned to the nearest gene versus distal genes aggregated across therapeutic areas. b. Proportion of prioritised genes that are distal to the GWAS lead variant (i.e. not the nearest gene) for four disease traits with high numbers of drug targets in ChEMBL, comparing Rosalind with Open Targets L2G predictions. c. Enrichment of prioritised genes for drug target-indication pairs from ChEMBL across clinical development phases. Points indicate odds ratios and horizontal bars indicate confidence intervals. d. Target capture performance comparison between Rosalind and L2G across clinical development phases. e. Overlap of clinically validated phase 4 drug targets recovered by Rosalind and L2G from GWAS loci. Venn diagram subsets are stratified by whether targets correspond to nearest genes or distal genes.

To assess whether these more distal predictions reflected biologically meaningful causal gene assignments, we tested enrichment of prioritised genes for known drug target-indication relationships across clinical development phases in ChEMBL. Rosalind predictions were enriched for established drug target-indication pairs, with enrichment increasing across later clinical stages (Fig. 2c). Importantly, this signal was not driven solely by nearest-gene predictions; Rosalind recovered targets of approved drugs that were missed by L2G (Fig. 2e), such as GIPR (e.g., Tirzepatide) and NDUFA7 (e.g., Metformin) for diabetes and GUCY1B1 (e.g., Riociguat) for hypertension.

### Osteoporosis case-study

To validate the utility of Rosalind in drug discovery, we applied it to predict the causal genes underlying variants associated with osteoporosis and experimentally tested these predictions in a disease-relevant functional assay. We focused on an estimated bone mineral density (eBMD) GWAS, a phenotype tightly linked to fracture risk and thus highly relevant to osteoporosis [20]. We took 1,103 conditionally independent genome-wide significant signals identified by Morris et al., 2019 [20] and for each used Rosalind to identify the plausible causal genes, extending the receptive field to 256kbp (Fig. 3a-b). This led to the identification of 239 high-confidence disease risk genes (DRGs) with the majority of them being distal (Fig. 3c). Not every variant yielded a gene prediction, which may be attributable to several factors: the lead variant may not be causal, the receptive field may not encompass the true causal gene, or limitations of the model. Encouragingly, several of the predicted DRGs are known to be causal in osteoporosis, including *TNFRSF11A* (RANK) and *TNFRSF11B* (OPG), which belong to the RANK/RANKL/OPG osteoclast pathway that is targeted by the anti-resorptive OP therapeutic, Denosomab (Anti-RANKL). Rosalind also predicted risk variants alter expression of *COL1A1, COL1A2*, and *WNT1*, which have been implicated in OP via whole exome sequencing (Fig. 3c).

**Fig. 3.**
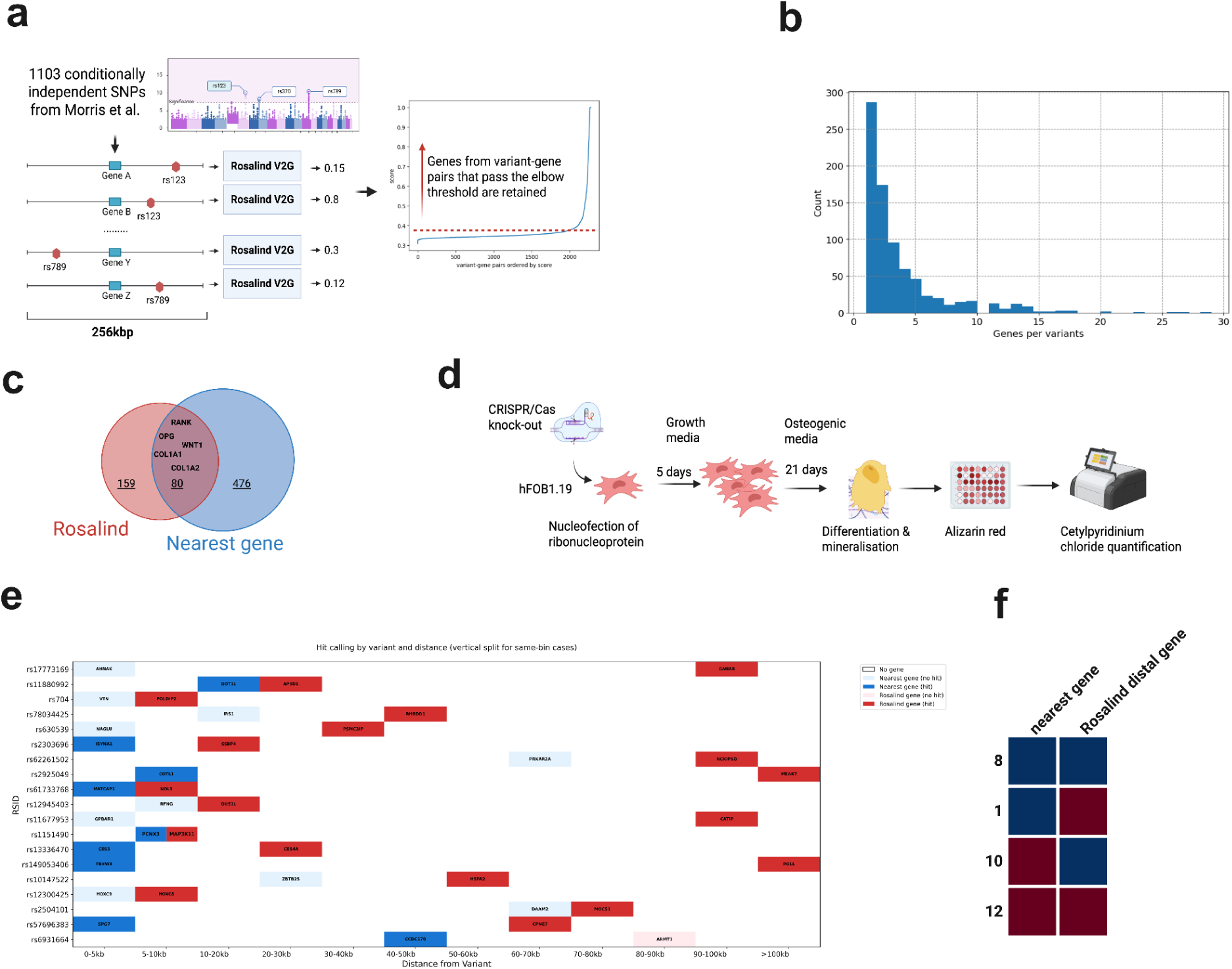
Using Rosalind for target discovery in Osteoporosis. a. All variant–gene pairs within a receptive field of 256kbp are scored by Rosalind. Genes above the elbow threshold are kept as candidate causal genes. b. The distribution of protein-coding genes around the 1103 lead variants from the eBMD GWAS (Morris et al. [20]) when using a window of 256kbp. c. Overlap of Rosalind-predicted genes with nearest genes is shown, with genes known to regulate eBMD highlighted. d. Mineralization assay experimental workflow: arrayed CRISPR/Cas9 knock-out screen of hFOB1.19 human osteoblasts. e. Plot shows SNP RSID-gene pairs, where at least one hit was observed in the osteoblast mineralisation assay. Hit calling was performed using Region of Practical Equivalence (ROPE) analysis and results show hits at ROPE 0.5 or greater (‘moderate’ effect size) f. Experimental validation summary, with assay “hits” represented in blue and “no effect” represented in red.

At the disease risk variant rs914615, Rosalind predicted that *FDPS* was the effector gene, rather than the nearest gene, *THBS3* [20], which has recently been shown to regulate osteogenesis [31]. Osteoclasts are sensitive to bisphosphonates, a class of bone anti-resorptives that inhibit *FDPS* leading to toxic metabolite accumulation [32]. Despite this central pharmacological role, *FDPS* has not been a commonly prioritised effector gene at BMD GWAS loci, however, the rs2297480 and rs11264359 genotypes are associated with differential responses to bisphosphonate treatment [33].

Furthermore, Rosalind mapped rs603424 to *CHUK* rather than the nearest gene, *PKD2L1* [20], independently validating a recent Mendelian Randomisation study [34]. Distal gene *CHUK* phosphorylates inhibitors of NF-κB, a known regulator of bone density [35]. While enrichment of pleiotropic eQTLs at enhancers and transcription factor binding sites is well documented [36], that is, a single fine-mapped or conditionally independent variant can regulate expression of multiple genes, without functional validation, it is not clear which gene is associated with the trait of interest.

### Osteoporosis variant-to-function mapping in osteoblasts

Large-scale eQTL mapping and functional enhancer perturbation studies have reported that a single fine-mapped or conditionally independent regulatory signal can be associated with expression changes in multiple nearby and distal eGenes (pleiotropic eQTLs), making it challenging to establish whether a single eGene or multiple eGenes, acting in the same effector cell or different effector cells, influences the disease trait of interest [22]. To test this, we used osteoporosis as an exemplar disease of common variation because core effector processes occur in a small number of well-characterised bone cell types for testing variant-to-function relationships: osteoclasts resorb damaged bone, osteoblasts synthesize new bone and osteocytes perform several functions, including regulating the balance of resorption and formation [21]. Given the practical limitations of designing phenotypic assays with disease-relevant endpoints for all three, we focused on human osteoblasts, a cell type that has been successfully targeted by bone anabolic therapies, but where therapeutic benefit is often only achieved for 12-24 months [37]. We developed an arrayed CRISPR/Cas9 knock-out screen to assess osteoblast mineralization, a key cellular endpoint that is relevant to bone formation (Fig. 3d). We confirmed the assay was fit-for-purpose by screening known regulators of osteoblast differentiation and mineralisation, such as those associated with osteogenesis imperfecta. To specifically assess Rosalind’s ability to predict distal causal genes, we conducted a side-by-side study. For each conditionally independent variant, we knocked out both the distal gene predicted by Rosalind and the nearest gene counterpart for the same original variant, measuring the effects on mineralization (Fig. 3e and Supplementary Table). The results revealed that the Rosalind-predicted distal genes were significantly more likely to elicit a positive effect in the osteoblast assay compared to the nearest gene (Fig. 3f, McNemar’s test, p-value = 0.011); supporting the idea that distal genes can be more informative for trait-related phenotypes. This is in agreement with a recent study of inflammatory bowel disease, where a single cell eQTL variant was more likely to colocalise if the conditionally independent or fine-mapped risk SNP lay in a distal enhancer rather than near the promoter of the target gene. At 12 loci, we did not identify any genes that regulate osteoblast mineralisation and at eight loci, multiple candidate target genes regulated osteoblast mineralisation (Fig. 3e-f).

### Rosalind predicts a causal role for osteoblast primary cilium structure in osteoporosis risk

Considering only the risk variants where the nearest gene was not a hit in the CRISPR/Cas knock out screen of osteoblast mineralisation but the Rosalind predicted distal gene was a hit, two of ten eGenes have roles in cilia structure and were hits decreasing mineralisation in the osteoblast assay (Fig. 3e). *CATIP* enables primary cilium formation by remodeling actin [38] and *GANAB* ensures correct folding of functional ciliary mechanosensors *PKD1/PKD2* and is present in osteoblast-derived extracellular vesicles [39,40]. A large body of in vitro and mouse model evidence supports a role for primary cilia in osteolineage mechanosensing and in BMP, WNT and Hedgehog-related signalling, and skeletal anomalies are evident in rare ciliopathies [41, 42, 43]. However, large BMD/osteoporosis GWAS have not consistently prioritised core ciliogenesis genes as a major determinant of BMD [20]. This could be due to the fact that many cilia genes are under evolutionary constraint and common variant GWAS require frequent variants with reasonably large effect sizes.

## Discussion

In summary, our work demonstrates that Rosalind, a DNA large language model, can accurately nominate causal disease risk genes from GWAS loci without relying on proximity-based assumptions, thereby overcoming a major limitation of conventional variant-to-gene mapping. Applied to OP, Rosalind prioritised hundreds of high-confidence effector genes, the majority of which were distal to the associated variants, and successfully recovered established therapeutic pathways while highlighting additional, previously underappreciated targets such as *CHUK*. Importantly, systematic functional interrogation in a disease-relevant osteoblast mineralisation assay showed that Rosalind-predicted distal genes were significantly more likely than nearest-gene candidates to influence bone formation. While we can not rule out the possibility that the nearest genes are more active in a different osteoporosis effector cell, these data provide experimental support for distal regulatory mechanisms at OP loci. Together, these findings establish DNA language modelling as a powerful framework for translating non-coding genetic associations into actionable biology, uncovering novel genes and mechanistic insights and accelerating target discovery for complex disease.

## Online Methods

### Rosalind

We extended the Enformer architecture by replacing the suite of relative positional encoding functions with Rotary Positional Embeddings (RoPE) integrated directly into the attention modules. This modification enables the model to generalize effectively to sequences longer than those encountered during training. The foundation model was trained on the original Enformer dataset, maintaining the same hyperparameters as the baseline Enformer model [15].

For fine-tuning, we encoded each variant by processing both the reference and alternate allele sequences through the pre-trained encoder, subsequently concatenating the resulting embeddings (Fig. 1 c). A lightweight discriminator was then appended to these representations and trained to distinguish causal from non-causal variants, directly aligning the model’s output with PIP-derived ground truth. Finally, as a form of data augmentation, we introduced non-causal variants at random positions within the input window during training to improve robustness.

### GeneGenie

GeneGenie is a deep neural network trained to predict discrete expression bins directly from promoter DNA sequence (–1 kbp to +100 bp relative to the transcription start site) by learning both local motif features and longer-range dependencies. To benchmark our fine-tuned self-supervised model against GeneGenie, we first re-trained the authors’ original model, which comprises a CNN, a transformer, and an LSTM, reproduced their results on the held-out test sets and obtained results for validation sets. Next, we trained our model on exactly the same promoter dataset the authors used, then obtained the identical validation and test splits reported in their manuscript and ran inference on each promoter sequence in those sets, assigning each to one of the same expression bins. Performance was measured in terms of PearsonR. We used both validation and held-out test data from GeneGenie’s original evaluation. Specifically, we are reporting results for the complex media, on the combined test set, which includes both, the random and native MPRA sequences.

Built on an Enformer backbone with a two-layer convolutional stem that downsamples 110bp to a 55bp representation before two transformer blocks, achieved 𝑅 = 0.98 under the same conditions on the test set. While we did not perform exhaustive ablations, we attribute this gain to more aggressive local feature aggregation via the simplified convolutional tower and to stable reverse-complement equivariance through data augmentation rather than parameter sharing [28].

### Osteoblast MPRA

We used a lentivirus-based massively parallel reporter assay (lentiMPRA) [27] for high-throughput functional characterization of candidate regulatory sequences (CRSs). Pools of 270-bp candidate enhancers or promoters are synthesized flanked by constant PCR handles; in a two-step PCR these are first fused to a minimal promoter and vector overhangs, then endowed with a 15-bp randomized barcode in the 5′ UTR of an EGFP reporter. The barcoded library is cloned into a lentiviral backbone, packaged into virus, and used to infect a human primary fetal osteoblast cell model [[KRT](https://www.cellapplications.com/human-osteoblasts-hob)], yielding stable “in-genome” integration of each CRS–barcode construct across millions of cells. Three replicates of infected cells are harvested: genomic DNA and total RNA are co-extracted, and barcode sequences are PCR-amplified (with unique molecular identifiers) to generate separate DNA- and RNA-seq libraries. Sequencing reads are first processed by the MPRAflow “association” utility to map barcodes back to their originating CRS then by the “count” utility to tally DNA and RNA barcode counts per element, normalize for library representation, and compute log₂(RNA/DNA) transcriptional activity for each CRS. The resulting per-element fold-change values provide a readout of the enhancer or promoter function across thousands of sequences.

### Osteoblast MPRA - Mutant generation strategy

Mutant candidate enhancers were generated based on positive control sequences active in the primary fetal osteoblasts extracted MPRA performed by Weiss et al. [[REF](https://elifesciences.org/articles/63713)]. A mutant generation strategy was defined by exhaustively “tiling” small contiguous mutation windows along a seed sequence with random selected bases to replace the original ones. For a sequence of length *s_length_* and a mutation window of length *mw_length_*, this strategy allows for the generation of up to *m_count_* mutants where *m_count_* = *s_length_* + 1 - *mw_length_*.

By using this strategy on positive control sequences (*s_length_* = 270) and *mw_length_* from 1 to 5, we generated 4,020 mutants based on the three enhancers driving the highest level of expression in the work of Weiss et al. Additional 7,518 mutants were generated starting from the next seven most active positive controls by allowing *mw_length_* from 1 to 4. This protocol resulted in a total of 11,538 mutant sequences.

### GTEx finemapped variant gene pairs

We obtained cis-eQTL summary statistics from the GTEx v8 release, encompassing 49 human tissues and nearly 17,000 samples, to serve as our input dataset. These summary statistics were then fine-mapped using the Sum of Single Effects (SuSiE) framework, which iteratively models multiple causal variants per locus and outputs posterior inclusion probabilities (PIPs) for each variant. In line with common practice, we classified variants with PIP > 0.9 as “causal,” reflecting high-confidence inclusion in credible sets and designated those with PIP < 0.01 as “non-causal,” effectively excluding them from credible models. To create a tissue-agnostic framework, we collapsed results across all tissues by taking the union of causal variants and non-causal variants, thereby formulating a binary classification task of whether a given variant influences gene expression in any tissue context. This approach reframes eQTL fine-mapping from a multi-tissue profiling problem into a single, global prediction of regulatory impact.

### Benchmarking Rosalind on held out GTEx finemapped variant pairs

We excluded a test set comprising cis-eQTLs from chromosomes 1 and 2, from model training and validation and used it to evaluate performance. The comparison focused on the tissue-agnostic classification of variant-gene pairs, using the same label definitions as those established during training. We constructed a balanced set where the closest non-causal variant was chosen as a matching negative for every causal variant in this set. Matched negatives were paired with the gene from the causal V2G pair to construct non-causal V2G pairs. We considered 1678 causal and 1586 non-causal V2G pairs for benchmarking, excluding variants for which an input sequence window could not be constructed for any of the predictors being compared due to restrictions on input DNA sequence length.

For Rosalind, predictions were generated as described above. For Enformer, we constructed reference and alternative sequence inputs by in silico insertion of the relevant allele, with the DNA sequence centred at the gene’s TSS in the V2G pair. We then compute scores for each output head as *log_2_-fold-change(alternate allele prediction/reference allele prediction)*. For CAGE predictions, the signal was aggregated within a 384 bp window centred on the gene TSS (3 prediction bins for Enformer). Scores were averaged across all CAGE output heads to produce a tissue-agnostic predictor. We took the absolute average (mean) log_2_-fold-change as the tissue-agnostic predictions. In addition, we considered two naive baseline predictors. The first predictor, called the nearest gene baseline, labels a V2G pair as causal if the variant’s nearest gene is the gene. The second predictor, called distance baseline, labels all V2G pairs where the distance between the variant and the gene falls within a defined threshold as causal.

We compared the predictive performance of these predictors on AUPRC and AUROC across different stratifications of the test set, including distance bin and whether the V2G gene is the closest gene to the variant. Stratification was performed after creating non-causal V2G pairs via the matching process described above. Confidence intervals were estimated by bootstrapping V2G pairs 500 times.

### Broad applicability of Rosalind for GWAS causal gene prediction

We ranked the traits in Open Targets by the number of genes with a completed phase 4 clinical trial, which typically indicates drug approval. For evaluating Rosalind, we selected four diseases from the top 20, spanning distinct therapy areas, and excluding cancer (due to somatic contributions) and neurological diseases (due to heterogeneous phenotype definitions and low number of genome-wide significant associations). For each trait (type 2 diabetes, hypertension, asthma and psoriasis), we selected the largest GWAS in Open Targets with confident SuSiE fine-mapping: FINNGEN_R12_T2D, FINNGEN_R12_I9_HYPTENS, FINNGEN_R12_J10_ASTHMACOPDKELA, FINNGEN_R12_L12_PSORIASIS. Across the four studies, we extracted 937 lead variants and scored all genes located within the models receptive field, resulting in 3766 V2G pairs. We compared against L2G scores for the same loci, made available in Open Targets. All specified data sets were obtained from Open Targets v25.03.

For downstream comparisons, we defined the L2G gene set as all genes with a score > 0.5, a commonly-used threshold in the community. To ensure comparable sizes, we defined a matching gene set from Rosalind predictions by selecting the top n ranked genes, where n equals the number of genes passing the L2G threshold. Enrichment analysis was performed using Fisher’s exact test and computing Wald confidence intervals on the resulting odds ratios.

### eBMD GWAS

We leveraged a large-scale genome-wide association study (GWAS) of estimated bone mineral density (eBMD) conducted by Morris et al. [20]. This study examined associations between genotype and heel quantitative ultrasound parameters, specifically, speed of sound (SOS) and broadband ultrasound attenuation (BUA), in 426,824 White British individuals from the UK Biobank [26]. The authors identified 1,103 conditionally independent, genome-wide significant SNPs (Supplementary Table 2 in [20]). For each lead variant, we defined the *nearest gene* as the protein-coding gene whose transcription start site lay closest to the variant position, and used this set of nearest genes as a baseline for comparison with Rosalind predictions.

Full methodological details are available in the original publication.

To link these variants to putative target genes, we identified all the windows centered on transcription start sites (TSS) of protein-coding genes that contain the lead variant.

This resulted in a comprehensive set of variant–gene pairs for downstream analysis.

We then applied our fine-tuned self-supervised model to predict variant-induced perturbations in regulatory activity for each gene locus. For genes appearing multiple times (due to overlapping windows), we retained the highest predicted score per gene. Finally, we ranked all genes by their maximal prediction score and identified a natural inflection point in the score distribution using the elbow method to define a high-confidence gene set for further experimental validation.

### Osteoblast CRISPR/Cas9 knockout arrayed screen of mineralisation

hFOB1.19 (ATCC) were cultured in flasks in DMEM/F-12, 10% FBS (Sigma F7524 lot.0001663052), antibiotic/antimycotic (Gibco 15240-062) and G418 sulfate (ab144261) at 34oC until 80-90% confluent. Cells (∼5x10^5) were nucleofected in a 96-well Nucleocuvette (Lonza) with ribonucleoprotein (Cas9, IDT and sgRNA, Synthego at 1:1.5) using the manufacturer’s protocol (Lonza; **DS 138** and solution **SE**). We used three sgRNAs per gene per well to ensure knock-out (Synthego). Cells were seeded at 25,000 cells per well in 48 well plates (12 plates per nucleofection), with CRISPR/Cas DNA cutting control guides (4 TRAC and 4 TNP2, neither of which are expressed in osteoblasts), spread equally over internal and edge locations. Each plate also included a negative mineralisation control (ALPL) and a positive mineralisation control (TP53). Cells were grown to 90-100% confluency for four days before inducing differentiation with osteogenic media (MEM Alpha, 10% FBS, antibiotic/antimycotic, G418 and 50 μg/ml ascorbic acid, 5 mM β-glycerophosphate) plus 100 nM Dexamethasone at 39oC (hFOb1.19 are transformed with a temperature sensitive SV40 construct). Media was changed every 2-3 days for 14 days and for the last seven days, the cells were cultured in osteogenic media alone. After 21 days, eight plates were fixed and stained with Alizarin red using standard methods and quantified using the cetylpyridinium chloride (CPC) quantification colorimetric (570 nm) plate reader (Varioskan Lux) assay. Remaining plates were banked for RNA isolation or DNA extraction to confirm the gene knock-out. The process was independently repeated three times. Outlier wells were removed with the Interquartile Range method (IQR multiplier of 2). A correlation matrix was computed across plates to identify outlier plates and the correlation between different nucleofections. Cutting controls (baseline) were averaged across the plate and plates were normalised to the average of all baseline controls to correct for plate variability. Assay quality was determined by comparing baseline controls with positive and negative controls using the strictly standardised mean difference (SSMD) metric [24]. We confirmed the absence of edge effects using a Bayesian linear model while taking into account plate effects. “Hit” calling was performed using a hierarchical Bayesian linear model while taking into account plate effects. We called hits when the region of practical equivalence score (ROPE) threshold >= 0.5 [25].

